# What happens to oak growth and survival when there is both competition *and* browsing?

**DOI:** 10.1101/2020.03.09.983437

**Authors:** Anna M. Jensen, Linda K. Petersson, Annika Felton, Magnus Löf, Maria Persson

## Abstract

Competition from neighboring vegetation and browsing by large herbivores are two of the most important factors affecting the structure and dynamics of temperate forests. While the previous literature has been able to identify individual negative effects from competition or browsing, no one has yet identified and quantified both the individual and the joint effects. Still, when plants face both competition and browsing, it is possible that the combined effect is not simply a sum of the individual negative effects, but perhaps a more complicated situation, where plants perform either better (in case there is also a facilitative effect from the neighboring vegetation) or worse (if the effects amplify each other) than they would if experiencing only one of the two factors. In this paper, we focus on regeneration of oak (*Quercus robur* L) to study these questions. We analyze a rich data set from a large long-term field experiment conducted at multiple sites in mixed temperate forests in southern Sweden over almost a decade. By the use of four separate treatments on each site – (i) neither competition, nor browsing, (ii) only competition, (iii) only browsing, and (iv) both competition and browsing – we can identify and quantify both the individual and combined effects of competition and browsing on oak growth and survival. We find that both competition and browsing individually affect growth and survival negatively. For growth, browsing has the largest effect, while competition is the larger problem from a survival point of view. When the plant experiences both competition and browsing, the combined, negative, effect is larger than either individual effect for survival, but for growth, the relationship is more complicated, and the surrounding woody vegetation offers at least some protection for the oak, reducing the negative effects from browsing.

## Introduction

Competition from neighboring vegetation and browsing by large herbivores are two of the most important factors affecting the structure and dynamics of temperate forests. Still, while the individual effects from competition or browsing are starting to be better understood [1–8], little is known empirically about their joint effects. In addition to implying competition for above- and below-ground resources, surrounding woody vegetation can have a facilitative effect by protecting the plant from browsing [9–14]. Thus, when plants face both competition from neighboring vegetation and browsing by large herbivores, it is possible that the combined effect is not simply a sum of the individual negative effects, but perhaps a more complicated outcome, where plants perform either better (in case there is indeed a facilitative effect) or worse (if the effects amplify each other) than they would if experiencing only one of the two factors. Understanding the individual and combined effects of competition and browsing is therefore interesting and highly relevant for policy and management. In this paper, we focus on oak regeneration to study these questions.

Oaks are important for ecosystem services in temperate forests. For example, they are one of the most critical tree genera for endangered invertebrates, lichens, fungi and birds, and in Sweden, more than 50% of all red-listed species on forest land depend on broadleaved forests often dominated by oaks [15–18]. Additionally, oaks provide wood production, recreation and other cultural services along with possibilities for adaptation of forest management to climate change due to their tolerance against various abiotic and biotic disturbances [19–21].

A widespread failure of natural oak regeneration has been cause for concern for over a century [22–25]. The reasons for this failure are still not well understood. Multiple factors probably contribute to poor oak regeneration, but changes in long-standing disturbance regimes (resulting in denser and darker forest stands that imply increased competition) and today’s high browsing pressure from wild ungulate browsers (deer and moose) are factors likely to affect regeneration success [6, 26–29]. However, it may be difficult to determine the actual causal relationship capable of explaining the negative regeneration responses when oaks face both competition and browsing. Is either competing vegetation or browsing individually the main cause for a reduction of the oak regeneration niche, or do these two factors interact to cause regeneration failures in oak?

The overall aim of the paper is to identify and quantify the individual and combined effects of competition and browsing on oak performance, i.e. growth and survival. To better understand the mechanisms through which competition and browsing affect oak performance, we (i) analyze determinants of how the neighboring vegetation community develops, as well as (ii) factors explaining the probability that browsing occurs. We then (iii) focus on the individual and combined effects of competition and browsing. Specifically, we ask the following questions. First, do oaks facing competition from neighboring vegetation (but no browsing) experience less growth and/or lower survival rates than oaks facing neither competition nor browsing? Second, do oaks facing browsing (but no competition) experience less growth and/or lower survival rates than oaks facing neither competition nor browsing? Third, how do the growth and survival outcomes for oaks facing both competition and browsing differ from oaks facing neither competition nor browsing, and how do they compare with oaks facing only competition or browsing?

To answer these questions, we analyze a rich data set from a large long-term field experiment conducted at multiple sites in mixed temperate forests in southern Sweden. The field experiment, focusing on the performance of English oak (*Quercus robur* L)., had at the last measurements run for almost a decade following establishment [13]. By the use of four separate treatments on each site – (i) neither competition, nor browsing, (ii) only competition, (iii) only browsing, and (iv) both competition and browsing – we can identify and quantify both the individual and combined effects of competition and browsing.

## Methods

### Sites and experimental design

The long-term field experiment was established in 2007 and conducted in ten mixed broadleaved forests across southern Sweden (Fig. 1). The 30-year mean annual precipitation decreases from about 1000 mm in the west to 500 mm in the east of the region, and the mean temperature ranges from −3°C in January to 16°C in July [30]. The sites are located 5-230 m above sea level. All sites were previously semi-open pastures or small fields, abandoned 50-90 years ago, and closed canopy mixed forest have subsequently developed through planting and secondary succession. Today all sites are of conservation interest with high biological values associated with large oaks in the overstory. Due to low rates of oak recruitment, conservation-oriented thinnings were conducted in the winter of 2002/2003 to facilitate oak regeneration. For further site and thinning description, see Götmark [31].

**Figure 1.**
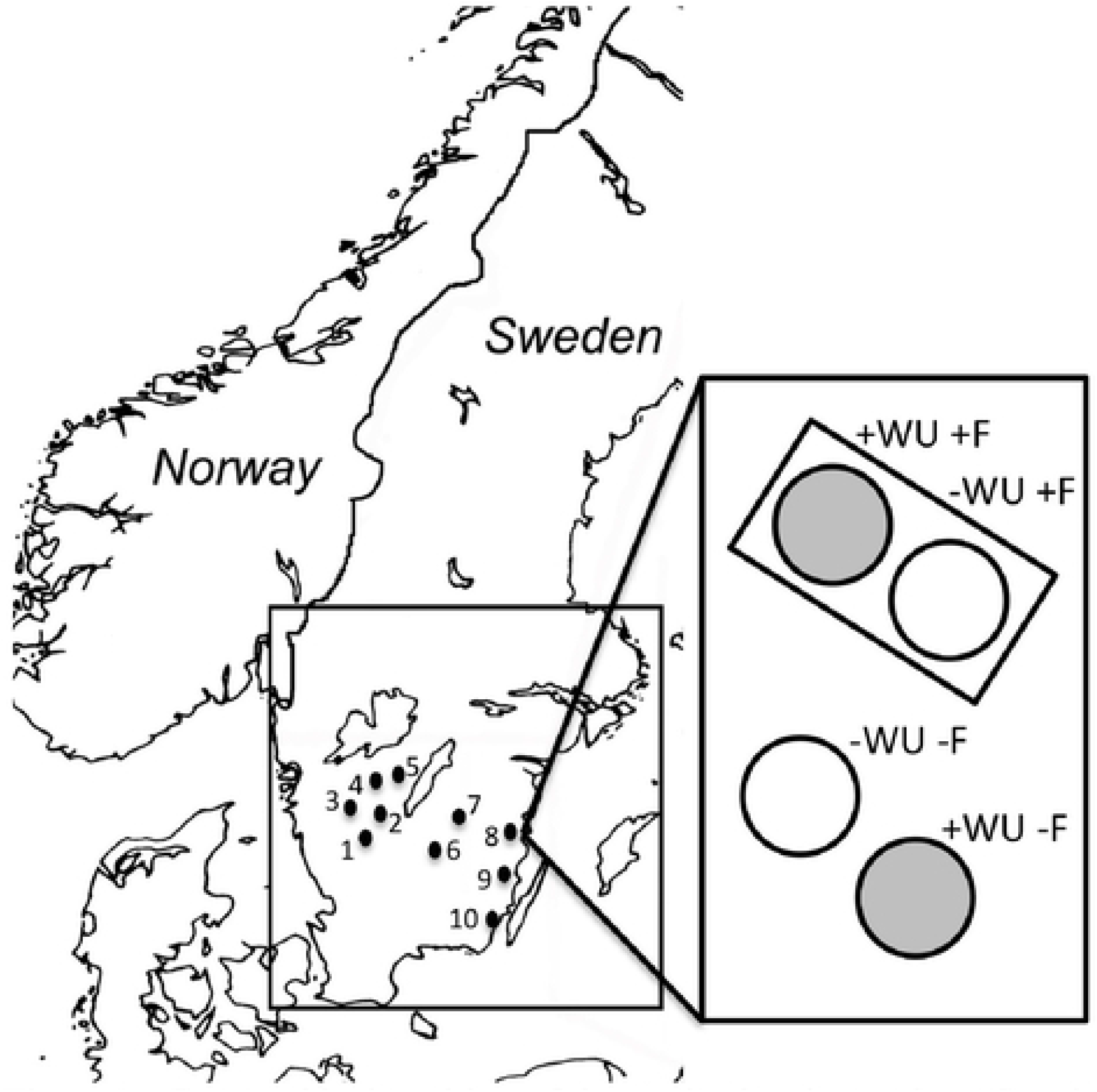
Geographical position of the study sites in southern Sweden and the experimental design. Mixed broadleaved forests used: (1) Rya åsar, (2) Sandviksås, (3) Östadkulle, (4) Karla, (5) Strakaskogen, (6) Norra Vi, (7) Aspenäs, (8) Fårbo, (9) Emsfors and (10) Lindö. The enlarged figure illustrates the experimental design at one site with four 30 m^2^ treatment plots; two with woody understory (gray circles, +WU) and two without woody understory (white circles, -WU). Two of the treatment plots were fenced (+F). [1-column figure]

The experiment was designed to test and quantify if naturally regenerated woody understory offers protection for young oaks against common ungulate browsers [see 13]. In December 2007 and January 2008, a total of 1200 bare-root oak (*Quercus robur* L.) saplings were planted in four 30 m^2^ plots at each site; two with naturally regenerated woody understory and two without (see enlargement in Fig. 1). Fences excluded moose (*Alces alces*), roe deer (*Capreolus capreolus*) and hare (*Lepus europaeus*) from two plots per site, generating four treatment combinations: fenced with woody understory (+WU+F); fenced without woody understory (-WU+F); not fenced with woody understory (+WU-F); and not fenced without woody understory (-WU-F) (Fig. 1). For more information, see Jensen et al. [13]. In order to maintain the -WU treatment over time, any sprouts of surrounding vegetation were manually cut down to <15 cm in height (between 2008-2011 every year and between 2012-2017 every second year). Five fence breaches occurred during the nine-year study. For each breach, the fence was mended within at most six months.

### Measurements and calculations

Data was collected during field campaigns (with a duration of two weeks) in April/May (2008, 2009, and 2010) and August/September (2008, 2009, 2013, 2015, and 2016). Oak survival was recorded, plants that were not found were assumed dead (oaks were individually marked and planted 1×1 m in the treatment plots, thus easy to relocate). The stretched height of the shoot (± 0.5 cm) was recorded for each oak, as well as the stem basal diameter (± 1 mm). The height to diameter (HD) ratio was calculated as the stretched height divided by the stem basal diameter. The percentage of herbivory was calculated as the proportion of oaks in a treatment plot damaged by browsing. When possible the herbivore responsible for the damage was identified based on Kullberg and Bergström [32] and classified as moose/roe deer (from here on referred to as ungulate), hare, or vole/mice.

Characteristics of the woody understory for each plot was determined around five randomly chosen planted oaks in each treatment plot by recording mean height (cm), stem density (stems m^-2^), and species composition. Broadleaved tree species that are highly preferred by ungulate browsers (naturally regenerated *Quercus robur, Quercus petraea*, *Populus tremula*, *Salix caprea and Sorbus aucuparia*) [32, 33] were grouped. Woody understory species were included if taller than 20 cm. Herbaceous characteristics in the treatment plots were determined by recording mean height (cm) and ground cover (%).

Light availability was estimated using hemispherical photographs in the fall of 2008, 2009, 2013, 2015 and 2016. One photograph was taken at 120 cm above ground level in each treatment plot using a Nikon Coolpix 8800 VR digital camera, with a LC-ER2 fisheye lens. The camera was kept horizontal using a spirit level and oriented against the magnetic north using a compass. Images were thresholded using SideLook version 1.1.01 [34] and then analyzed with software Gap Light Analyzer to calculate leaf area index (LAI) [35]. The relationship between LAI and relative light availability is shown in figure S2 (light data is available on request).

We calculated an index for the relative effect of competition (RCI), browsing (RBI) and the combination of competition and browsing (RCBI) as follows:

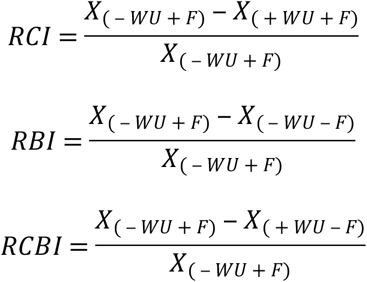

where X is individual performance variables (survival, basal diameter, height and HD ratio) for the respective treatment combination [36].

### Regression analysis

In addition to the descriptive analysis, we also perform regression analysis to dig deeper into the relationship between variables. When analyzing factors that could potentially explain the growth of the oaks, we use linear regression models. Because there is a great deal of unobserved heterogeneity between the sites and over time, we use the fixed effects (i.e. within) estimator, rather than OLS (ordinary least squares). Specifically, we include fixed effects for every individual combination of site and field campaign (i.e. a site by time fixed effect). Briefly, these fixed effects will capture unobserved heterogeneity that varies between sites (such as soil quality), between years (such as precipitation patterns) and between each individual combination of year and site (such as the emergence over time of a large moose population in one particular site). We therefore drastically reduce the risk of biased results due to a misspecified model (omitted variable bias). Note that the rho values reported in the results tables indicate how much of the variation that is captured by the fixed effects. With rho values as high as 0.53, failing to control for the unobserved heterogeneity would imply running a serious risk of getting biased results.

We also analyze how the probability of browsing is affected by various factors. Since the dependent variable is then binary, we use logistic (i.e. logit) regressions. We again include fixed effects for every combination of site and field campaign. While the logistic regressions have the advantage of always producing predicted values that lie within the proper range, we also use the linear probability model as a robustness check. While potentially producing predicted values outside the 0-1 range, this model has the advantages of both being more easily interpreted and producing rho values which allows us to gauge the importance of controlling for unobserved heterogeneity.

Lastly, we are also interested in analyzing how various factors affect the oaks’ survival. Since we have data that is distinctly discrete by nature – the observations are grouped into (roughly) bi-yearly intervals – we use a survival model designed for discrete-time data. We focus on the logit model where we can both include fixed effects without problems and do not have to make the restrictive assumption about proportional hazards. We note that left-censoring of the data is not an issue here, since we know the exact time when the oaks were planted, and right-censoring is of course not a problem with a properly specified duration model. We specify the baseline hazard in the most flexible way possible by using duration-time specific dummies. Please note that this choice also solves the potential problem that the time between the various field campaigns may not be of exactly the same length. Due to the specification of the baseline hazard, we use site (rather than site by time) fixed effects for the survival analysis.

Stata 15 has been used for all regressions.

## Results and discussion

### Main results in brief

We find that both competition and browsing individually affect growth and survival negatively. For growth, browsing has the largest effect, while competition is the larger problem from a survival point of view. When the plant experiences both competition and browsing, the combined, negative, effect is larger than either individual effect for survival, but for growth, the relationship is more complicated, and the surrounding woody vegetation offers at least some protection for the oak, reducing the negative effects from browsing.

The overall aim of the paper is to identify and quantify the individual and combined effects of competition and browsing on oak survival and growth. Before engaging in that analysis, we will briefly discuss how the neighboring vegetation develops over time in our data, and then analyze the determinants of the probability of the oak being browsed.

### Development of the neighboring vegetation

The exclusion of browsing animals through fencing does not have a uniform effect on woody understory community structures. While mean height and density were unaffected by the fence, the species composition changes (Fig. 2A-D and Table S1), surprisingly reducing the proportion of browser preferred species (*S*. *aucuparia*, *P*. *tremula*, *S*. *caprea* and naturally regenerated *Q*. *robur* and *Q. petraea*) inside fences. This finding supports previous research which has demonstrated that browsing ungulates can have far-reaching effects on forest understory structure and composition [8, 37, 38]. Furthermore, the presence of woody understory vegetation negatively affected the occurrence of naturally regenerated oak, reduced the herbaceous cover and the light availability (i.e. greater LAI) within the treatment plot (Fig. 2D-F and Table S1). Although the age and time of establishment of naturally regenerated individuals was not recorded in the present study, it is possible that new establishment of browser preferred species was limited inside fences by for example shading by the surrounding vegetation (Table S2) [13, 39], and by increased seed predation from mice [40, 41]. For example, high local densities of moose can lead to increased light availability for understory plants, leading to lower levels of light competition, particularly in sites of relatively high productivity [42]. Boulanger et al. [38] reported that plant communities were more light-demanding outside than inside 10-year-old ungulate exclosures. Regardless of whether the browser preferred species were recruited before or after the beginning of the field experiment, all species grouped as browser preferred in the current study are light demanding [43–46], which may have affected their capacity to compete within a woody understory, especially as the light availability is limited also from the overstory trees.

**Figure 2.**
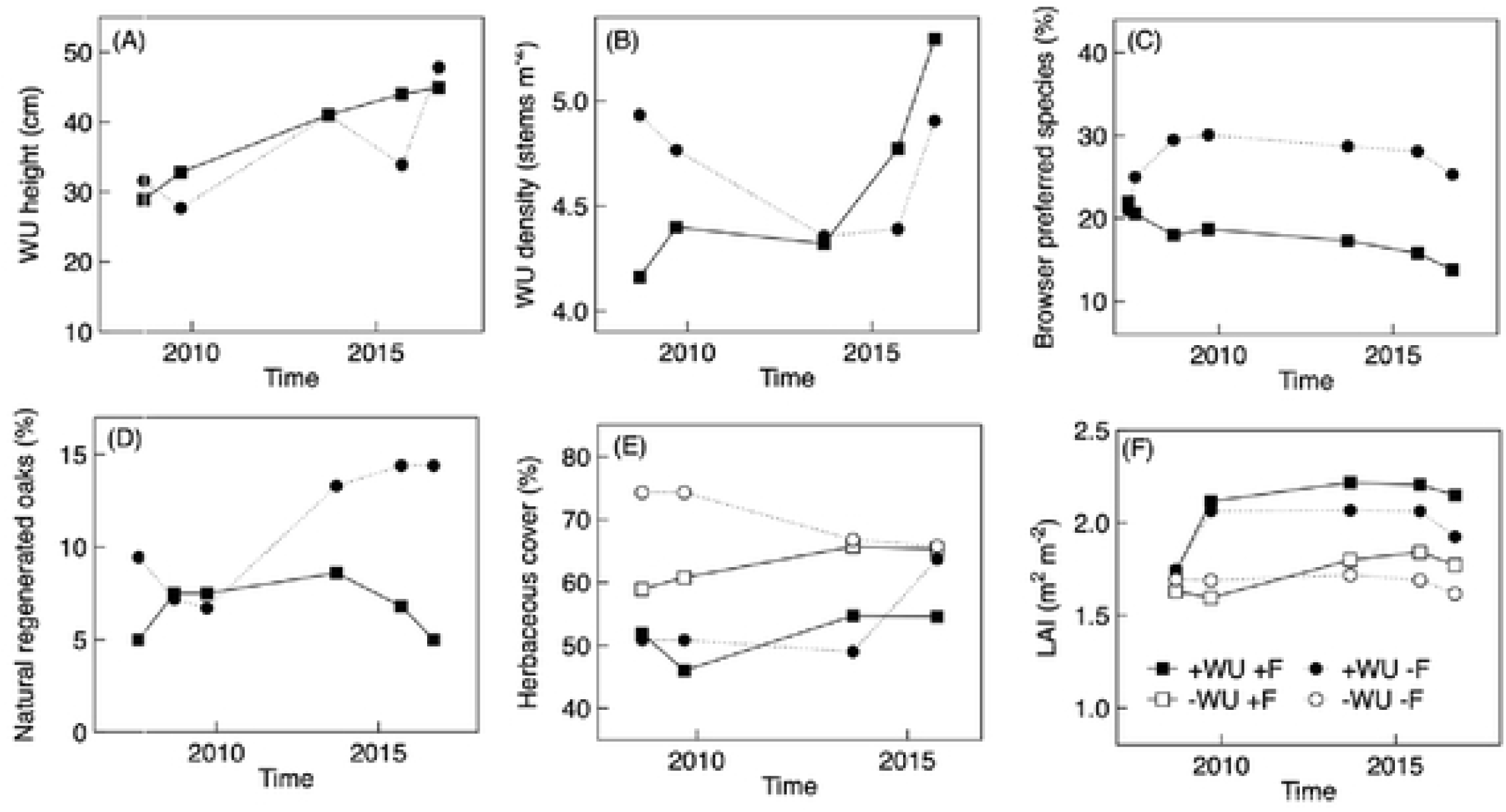
Development of (A-D) the surrounding woody understory (WU), (E) the herbaceous communities, and (F) light availability surrounding oaks grown under four treatment combinations in southern Sweden from 2007 to 2016. In panel A-D, only treatments with woody understory vegetation are shown. Browser preferred species (% of all stems, C) includes; *Sorbus aucuparia*, natural regenerated *Quercus robur* and *Q. pelraea, Populus Iremula* and *Salix caprea.* Whereas natural regenerated oaks (% of all stems, C) includes; *Q. robur* and *Q. petraea.* Mean values, n = 10. [2-column figure]

### Browsing probability

Earlier output from this field experiment has shown that the woody understory has a protective effect, reducing browsing on young oaks [13]. Our aim here is to test empirically whether this protective effect persists over time, but also to learn more about which characteristics of the woody understory and the oaks themselves that affect browsing.

We find that fencing provides better protection against browsing than growing the oak hidden in woody understory: Although we had a total of five fence breaches over the nine years only a single oak protected by a fence was browsed, while for oaks outside the fence, 9-67 % of all oaks were browsed (Fig. 3). However, while less effective than fencing, woody understory surrounding the oak plant still offers protection. We find that the probability that the oak plant will be browsed is highly significantly decreased in the presence of woody understory (Table 1), and the share of oaks that have been browsed is 5 to 30 percentage points lower in plots with woody understory (Fig. 3). These findings are consistent with previous short-term studies [e.g. 9, 11, 12, 14], while the present study demonstrated that neighboring vegetation could offer some protection from deer browsing over the long-term as well. It is worth noticing that we observe this protective effect even though the protective effect of neighboring vegetation is known to decrease with increased browser densities [9, 11, 47], and there are indeed large ungulate browser populations in southern Sweden (our study area) [48, 49]. From an ecological and management perspective it would be valuable to identify browser-density thresholds at which this protective effect disappears.

**Figure 3.**
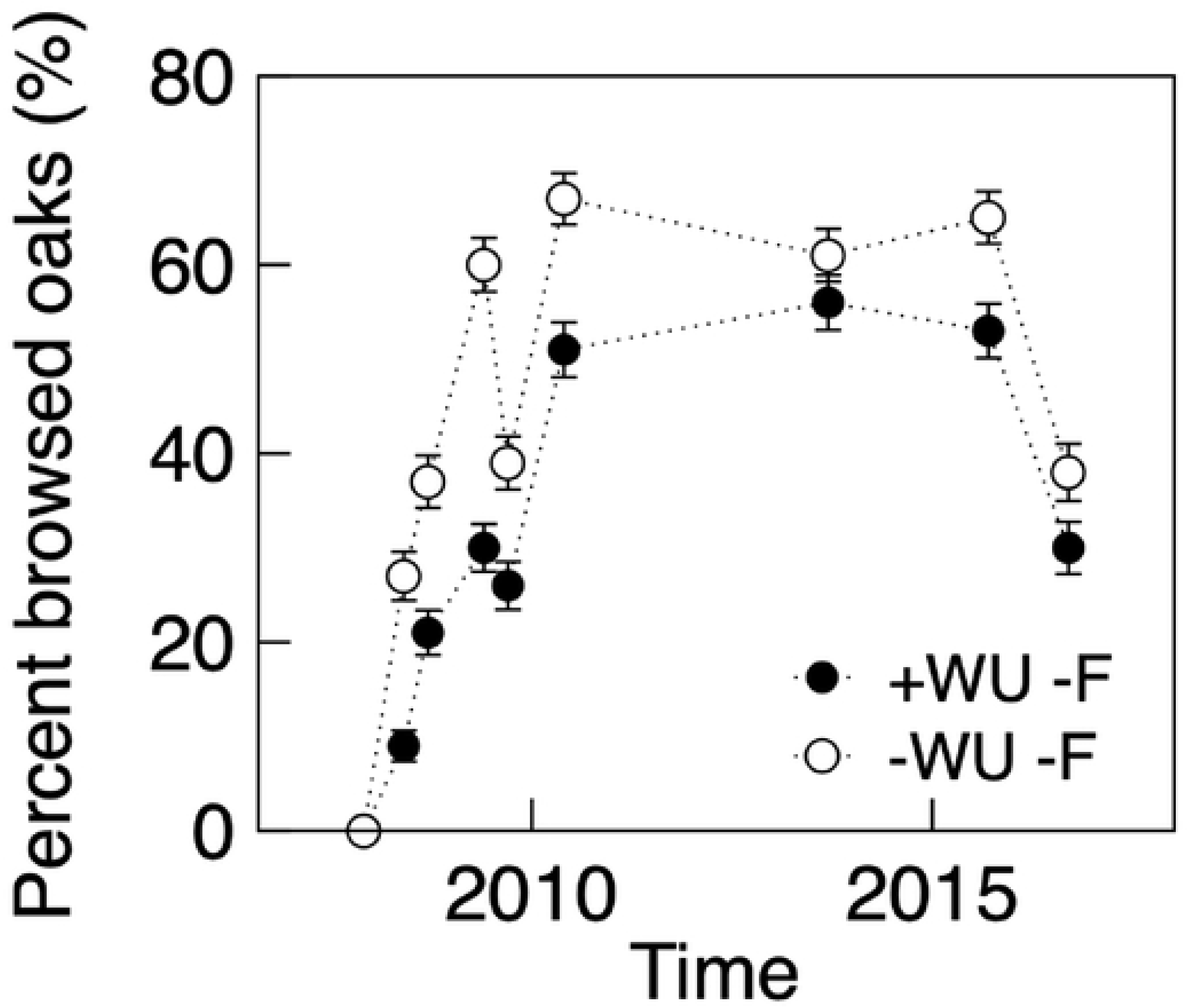
Percent oaks browsed in a treatment plot by ungulates on oaks grown in the presence (+WU, filled circles) and absence (-WU, open circles) of woody understory, both treatments without fence (-F). Mean values ± ISE, n = 10. [1-column figure]

**Table 1.** Regression results. How is browsing probility affected by woody understory and the oak’s own characteristics?

| VARIABLES | (1)<br>FE logit | (2)<br>FE logit | (3)<br>FE logit | (4)<br>FE logit | (5)<br>FE linear |
| --- | --- | --- | --- | --- | --- |
| Treatment: Woody understory, no fence | -1.020***<br>[0.000] | -0.839***<br>[0.000] | -0.868***<br>[0.000] | -1.123***<br>[0.000] | -0.189***<br>[0.000] |
| Oak was browsed in latest period |  | 1.329***<br>[0.000] | 1.335***<br>[0.000] | 1.530***<br>[0.000] | 0.313***<br>[0.000] |
| Oak's height in latest period (cm) |  |  | 0.004<br>[0.224] | 0.003<br>[0.449] | 0.000<br>[0.510] |
| Oak's basal diameter in latest period (mm) |  |  |  | 0.056*<br>[0.043] | 0.008<br>[0.060] |
| Constant |  |  |  |  | 0.326***<br>[0.000] |
| Observations | 2,857 | 2,658 | 2,654 | 1,581 | 1,612 |
| Pseudo R-squared | 0.0451 | 0.0918 | 0.0931 | 0.120 |  |
| Chi-squared | 131.9 | 250.1 | 253.2 | 195.5 |  |
| Adjusted R-squared |  |  |  |  | 0.113 |
| F-value |  |  |  |  | 59.48 |
| rho |  |  |  |  | 0.193 |

Notes: The dependent variable is an indicator variable taking the value 1 if the oak plant has been browsed in the current period, and 0 if not. The models are run on all observations for the treatments without fence. In columns 1–4, a logistic (logit) regression is run to assess the probability of browsing, adding fixed effects for every combination of site and field campaign. In column 5 (regression 5), we instead use the fixed effects (i.e. within) estimator to run a linear probability model. The fixed effects in column 5 still capture every combination of site and field campaign. *P*-values in brackets. \*\*\* $p < 0.001$ , \*\* $p < 0.01$ , \* $p < 0.05$ .

In terms of the oak’s own characteristics, the best indicator of whether an oak plant will be browsed in the current period is whether it was browsed in the last period: browsing in the previous period highly significantly increases the probability of browsing (Table 1). Similar findings have been reported for several other tree species, including important staple food species [50, 51]. Such re-browsing might be caused by increased nutrient concentration, or reduced concentration of plant secondary compounds, following browsing [51 and references therein]. However, other characteristics of the oak plant seem to matter less. We find no evidence that the height of the plant has an effect on the browsing probability (Table 1). As the average height of oaks outside fences were 50-75 cm at the end of the experiment, it is possible that they were not tall enough for the height to have a significant impact. There is limited support for the hypothesis that the basal diameter – acting as an indicator of canopy size [52] – matters. The basal diameter in the previous period has a positive coefficient and is significant when we estimate with the theoretically more appropriate logistic model (Table 1). However, since it is only significant at the 5 % level, and we do not get a significant coefficient when we use a linear probability model as a robustness check, we do not want to oversell this finding. We have also tried separating the observations, so that the models are run separately on the observations in the treatments with and without woody understory, and combined but allowing the effects of the oak’s characteristics to differ depending on treatment through the inclusion of interaction effects (results available upon request). The conclusions we draw are the same, indicating that while the treatment (i.e. the availability of woody understory) in itself affects the probability of browsing, it does not change the way the oak’s own characteristics affects browsing probability.

The character and structure of the individual woody understory community influences the probability of browsing. Growing in a plot with a high proportion of browser palatable species, or high herbaceous ground cover increased the risk of an oak being browsed, while a high surrounding woody vegetation decreases the risk. We find no evidence that the density of the woody understory or LAI affects the probability of oaks being browsed (Table 2). It seems likely that taller understory vegetation is more efficient at visually blocking the oak plants and thereby reduce the likelihood of them being detected by browsers [cf. 53–55]. While shrub communities with a high proportion of palatable species are more attractive to browsers, the risk of an individual plant being browsed might be dependent on the plant’s relative palatability within the community. These different scales of forage selection could explain the contradictory reports as to whether palatable neighbors increase [50, 56, 57], decrease [47] or has no effect [58] on the browsing probability of the focal plant. As oaks are highly palatable [32], it would seem they are at disadvantage at both these forage selection levels.

**Table 2.**
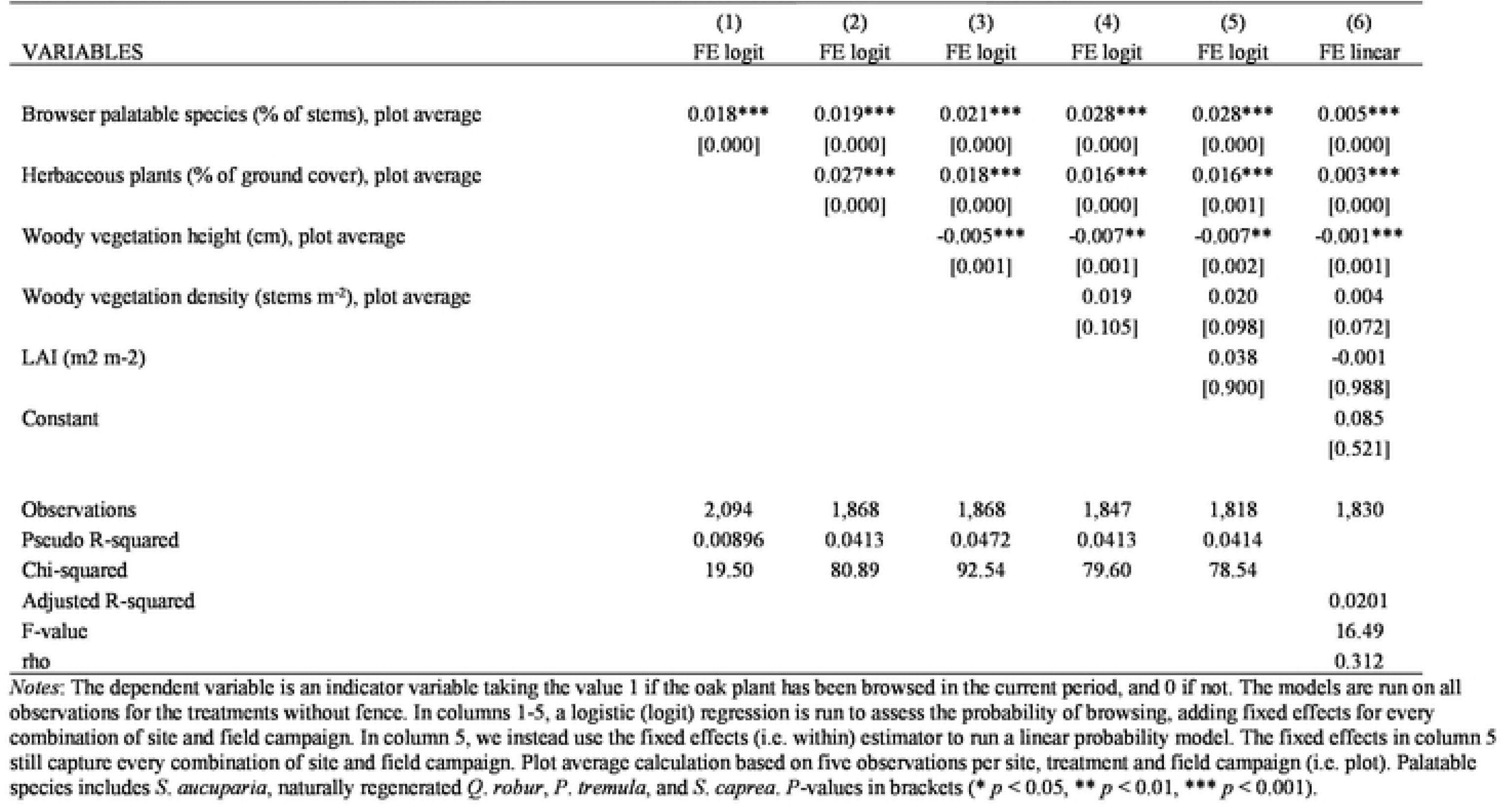
Regression results: The effect of woody understory characteristics on ungulate browsing probability on oak.

| VARIABLES | (1)<br>FE logit | (2)<br>FE logit | (3)<br>FE logit | (4)<br>FE logit | (5)<br>FE logit | (6)<br>FE linear |
| --- | --- | --- | --- | --- | --- | --- |
| Browser palatable species (% of stems), plot average | 0.018***<br>[0.000] | 0.019***<br>[0.000] | 0.021***<br>[0.000] | 0.028***<br>[0.000] | 0.028***<br>[0.000] | 0.005***<br>[0.000] |
| Herbaceous plants (% of ground cover), plot average |  | 0.027***<br>[0.000] | 0.018***<br>[0.000] | 0.016***<br>[0.000] | 0.016***<br>[0.001] | 0.003***<br>[0.000] |
| Woody vegetation height (cm), plot average |  |  | -0.005***<br>[0.001] | -0.007**<br>[0.001] | -0.007**<br>[0.002] | -0.001***<br>[0.001] |
| Woody vegetation density (stems m<sup>-2</sup>), plot average |  |  |  | 0.019<br>[0.105] | 0.020<br>[0.098] | 0.004<br>[0.072] |
| LAI (m<sup>2</sup> m<sup>-2</sup>) |  |  |  |  | 0.038<br>[0.900] | -0.001<br>[0.988] |
| Constant |  |  |  |  |  | 0.085<br>[0.521] |
| Observations | 2,094 | 1,868 | 1,868 | 1,847 | 1,818 | 1,830 |
| Pseudo R-squared | 0.00896 | 0.0413 | 0.0472 | 0.0413 | 0.0414 |  |
| Chi-squared | 19.50 | 80.89 | 92.54 | 79.60 | 78.54 |  |
| Adjusted R-squared |  |  |  |  |  | 0.0201 |
| F-value |  |  |  |  |  | 16.49 |
| rho |  |  |  |  |  | 0.312 |

Notes: The dependent variable is an indicator variable taking the value 1 if the oak plant has been browsed in the current period, and 0 if not. The models are run on all observations for the treatments without fence. In columns 1-5, a logistic (logit) regression is run to assess the probability of browsing, adding fixed effects for every combination of site and field campaign. In column 5, we instead use the fixed effects (i.e. within) estimator to run a linear probability model. The fixed effects in column 5 still capture every combination of site and field campaign. Plot average calculation based on five observations per site, treatment and field campaign (i.e. plot). Palatable species includes *S. aucuparia*, naturally regenerated *Q. robur*, *P. tremula*, and *S. caprea*. P-values in brackets (\* $p < 0.05$ , \*\* $p < 0.01$ , \*\*\* $p < 0.001$ ).

### Competition versus browsing: How do they affect growth and survival?

Having established that our various treatments do tend to affect both characteristics of the neighboring microhabitat and the probability of browsing, we now turn to identifying and quantifying the individual and combined effects of competition and browsing on long-term oak growth and survival.

### Competition effects on growth

Values for the index for the relative effect of competition (RCI) on oak height and basal diameter growth were positive the first year, indicating a facilitation effect, but decreased to - 0.20 and −0.33 by 2016 (Fig. 4A,B). Our regression analysis shows a statistically significant competition effect: Oaks grown in a treatment with woody understory and fence are significantly lower and have a smaller basal diameter than oaks grown in the reference treatment (no woody understory, but fence) (Fig. 4A,B, Fig. S1A,B, and Table 3). Our results are consistent with previous research that have reported both positive [7, 13] and negative [52, 59] effects on oak growth from woody understory. Analyzing the (HD) ratio, competition results in a statistically significant positive effect, meaning that the oaks prioritize height growth over basal diameter growth when facing competition (Fig. 4C, Table 3). The effects of woody understory competition on the HD ratio are consistent with previous research [5, 7, 59]. In an open-field experiment located in southern Sweden, Jensen and Löf [7] found that as woody vegetation overtopped oaks (*Q. robur*) it caused growth stagnation and a marked drop in survival of the oaks the following years. Furthermore, they found a great increase in the HD ratio of oaks growing among woody vegetation, consistent with the results in the present study. These findings suggest that in response to woody understory competition, the oaks alter their allocation patterns to increase light capture, but over time they do not have enough resources to keep up with the competitors.

**Figure 4.**
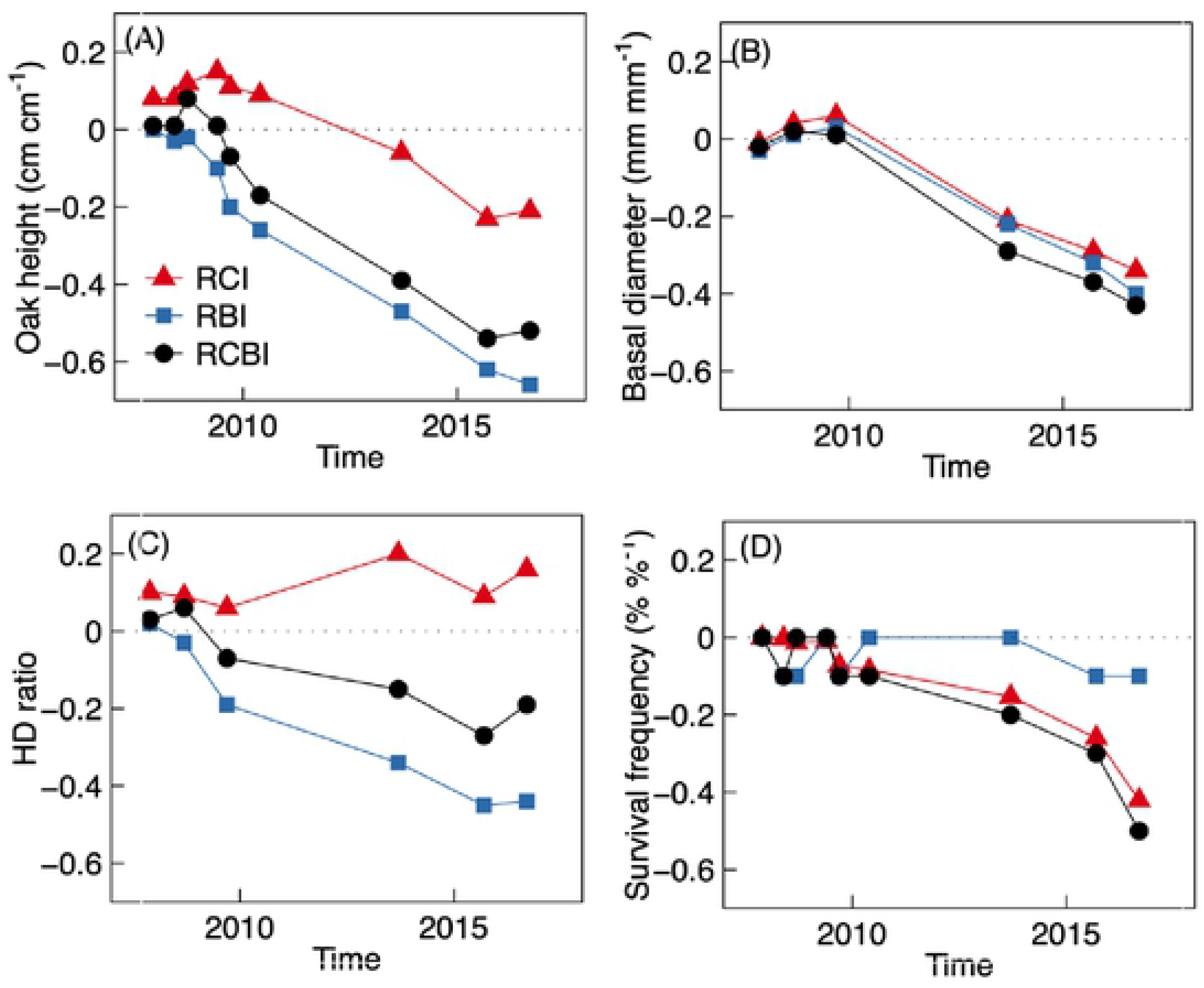
Relative effects of competition (the relative competition index (RCI), red triangles), browsing (the relative browsing index (RBI), blue squares) and the combination of competition and browsing (the relative competition and browsing index (RCBI), black circles) over time on oak (A) height, (B) basal diameter, (C) height to diameter (HD) ratio, and (D) survival. Positive values indicates a facilitation effect. Values are means (n = 10). [2-column figure]

**Table 3.** Regression results: Individual and combined effects of competition and browsing on oak growth and suvival.

| VARIABLES | (1) | (2) | (3) | (4) |
| --- | --- | --- | --- | --- |
|  | Total height (cm).<br>Linear FE | Basal diameter (mm).<br>Linear FE | HD ratio (cm cm<sup>-1</sup>).<br>Linear FE | Probability of death.<br>FE logit |
| Treatment: Woody understory, fence (competition effect) | -4.613***<br>[0.000] | -2.183***<br>[0.000] | 0.065***<br>[0.000] | 1.328***<br>[0.000] |
| Treatment: No woody understory, no fence (browsing effect) | -26.745***<br>[0.000] | -2.680***<br>[0.000] | -0.193***<br>[0.000] | 1.053***<br>[0.000] |
| Treatment: Woody understory, no fence (combined competition and browsing) | -18.860***<br>[0.000] | -2.891***<br>[0.000] | -0.083***<br>[0.000] | 1.759***<br>[0.000] |
| Constant | 73.472***<br>[0.000] | 12.550***<br>[0.000] | 0.696***<br>[0.000] |  |
| Observations | 7,830 | 4,489 | 4,487 | 7,939 |
| Adjusted R-squared | 0.139 | 0.0891 | 0.174 |  |
| F-value | 449.1 | 163.3 | 331.2 |  |
| rho | 0.456 | 0.346 | 0.220 |  |
| Pseudo R-squared |  |  |  | 0.330 |
| Chi-squared |  |  |  | 1305 |

Notes: For columns 1-3: Linear regressions estimating the effects of the various treatments on (1) oak total height in cm, (2) oak basal diameter in mm, and (3) oak HD ratio (cm cm<sup>-1</sup>). Models are estimated with the fixed effects (i.e. within) estimator, with fixed effects for every combination of site and field campaign. The reference group in the regressions is the treatment with neither competition, nor browsing (i.e. without woody understory, but with fence). Thus, the coefficients should be interpreted as the difference between the oaks planted in the other treatments, and oaks planted without woody understory, but with fence. Column 4: Discrete-time logistic (i.e. logit) duration model, analyzing the effect of various treatments on oak survival. The baseline hazard has been specified in the most flexible manner, with one dummy for each spell year. Note that a positive coefficient implies that the variable increases the hazard of dying. For the survival analyses, fixed effects are included for every individual site. For all columns: *P*-values in brackets (\* *p* < 0.05, \*\* *p* < 0.01, \*\*\* *p* < 0.001).

### Browsing effects on growth

Browsing strongly and highly significantly affects height development, basal diameter, and the HD ratio (Fig. 4A-C, S1A-C and Table 3). Values of the index for the relative effect of browsing (RBI) range between −0.03 - -0.66, 0.03 - -0.40, and 0.02 - -0.45 for oak height, basal diameter and HD ratio, respectively. In fact, our results show that browsing is a more important obstacle for growth than competition. As can be deduced from Table 3, oaks grown in the reference treatment with neither competition nor browsing, are on average 73 cm tall. Oaks grown in the treatment with only browsing are almost 27 cm shorter, while the oaks grown with only competition are only about 5 cm shorter (N.B. the coefficients show the difference against the reference treatment with neither competition, nor browsing, but we have also confirmed that there is a highly statistically significant difference between the treatments with only browsing and only competition, respectively. Results available upon request). We draw the same conclusions when looking at basal diameter: While competition is damaging for basal diameter growth, the effect of browsing is (significantly) more damaging. These findings are consistent with previous research that have reported negative effects on oak performance from ungulate browsing [e.g. 6, 60, 61].

Focusing on the HD ratio, we find an interesting result, where – again comparing against oaks grown with neither competition, nor browsing – oaks facing only competition prioritize height growth (i.e. there is a positive effect on the HD ratio) (Fig. 4C, Fig. S1C, Table 3). Browsing also has a significant (negative) effect on the HD ratio. In this case, it is more complicated to interpret this as saying anything about the plant’s growth strategy, since height loss due to browsing will in itself affect the HD ratio.

### Combined effects of competition and browsing on growth

Having established that competition as well as browsing have individual negative effects on oak growth, and that the latter effect is significantly stronger, we now ask what happens if there is both competition and browsing. For the oak’s height, we find that there is a smaller negative effect from facing both competition and browsing, than from facing only browsing (Fig. 4A, Fig. S1A, Table 3). Thus, while our results show a negative height effect from either competition or browsing, the surrounding woody understory that competes with the oaks also offers protection from browsing. In other words, when it comes to oak height, we find a competitive as well as a facilitating effect from the woody understory. When it comes to basal diameter, the story is different. Here, while, as outlined above, both competition and browsing have a negative effect on the basal diameter, it is actually even worse for the plant if there is both competition and browsing: The oaks grown in a woody understory outside the fence are even thinner than the oaks grown with only competition or browsing (all differences are highly statistically significant) (Fig. 4B, Fig. S1B, Table 3). Thus, overall our conclusion is that (i) competition is hurtful for both height growth and basal diameter growth, and (ii) browsing is even more hurtful for both. However, we also find that while plants facing browsing can use the protection of the surrounding woody understory to grow taller than if they only faced browsing, the plants do not have enough resources to use this facilitating protection to also grow wider. Thus, (iii) the combined effect of competition and browsing is especially hurtful for basal diameter growth, while the negative browsing effects on the oaks’ height is mitigated somewhat from the protection of the (competing) woody understory.

### Competition effects on survival

The first year after planting, survival was 98%, irrespective of treatment. Thereafter, the survival dropped in all treatment plots. For the oaks facing competition from surrounding woody understory, but no browsing, survival rates starts to drop already in the second year after planting, and over time there is a sizeable effect from competition on survival (Fig. 4D, S1D). Values of RCI increase from 0.00 to −0.4 (Fig. 4D). Focusing on our regression results, we also find clear indications that competition affects survival. If the oaks grow up facing competition from surrounding woody vegetation, but under the protection of a fence that prevents browsing, the hazard of dying is highly statistically significantly larger than if the oaks face neither competition, nor browsing (Table 3). This is in line with previous research (e.g. 5, 7, 59), and expected for a pioneer species like Q. *robur*. Many members of genus *Quercus* are intolerant of shade, and growth is often limited when available understory light drops below 20 % of the above canopy light levels [e.g. 43, 62] or when leaf area index (LAI) is around 2.9. In many forest understories, light rarely reaches above these levels, and oak survival thus benefits from partial thinning that reduces overstory competition [28, 59, 63]. However, partial thinning may complicate matters by also promoting competing understory woody vegetation [39, 43] i.e. enhancing completion pressure.

### Browsing effects on survival

Our results suggest that browsing affects survival less than competition. As seen in Fig. 4D, the difference in survival between oaks grown with neither competition, nor browsing, and those facing browsing is smaller than that found for oaks facing only competition. Values of RBI range from 0.00 to −0.01 (Fig. 4D, Fig. S1D). However, while smaller, our regression analysis still finds that the effect from browsing is highly statistically significant (Table 3). These results are in line with the literature which has established that, although oaks are preferred by browsers, they can survive moderate browsing pressure for extended periods, provided there is enough available light (i.e. low competition pressure), and they are therefore considered tolerant to browsing [25, 39, 64]. With values of LAI ranging from 1.6 to 1.7 in the treatment with only browsing in the present study, it is possible that repeated browsing, which depletes the oaks’ resources over time, caused the mortality observed in this treatment [65]. Furthermore, previous studies have demonstrated that ungulate browsing can severely limit oak growth [6, 32, 61] and that high deer densities may limit recruitment of palatable tree species [3, 48, 60]. Our findings further support these previous observations, as we could demonstrate that browsing had a greater negative effect than competition on oak growth, but that the reverse relationship was true for oak survival (Table 3).

### Combined effects of competition and browsing on survival

Having found that competition and browsing both have an individual negative effect on the survival of oaks, we now ask what happens when oaks face both competition and browsing. In our regression analysis, we find that oaks facing both competition and browsing have a highly significantly larger hazard of dying than plants in the reference treatment (neither competition, nor browsing) (Table 3). Looking at the RCBI (index of the relative effect of both browsing and competition) values, we find that the risk of oak plants not surviving is even greater for the oaks facing both competition and browsing, than for oaks facing only competition or browsing (Fig. 4D, Fig. S1D). In other words, unlike for growth, where the surrounding woody vegetation implied competition, but also some protection from browsing, here we do not find any facilitation mechanism, but the combined effect of competition and browsing on survival is even greater than either effect individually.

## CONCLUSIONS

### Main conclusions

This paper is the first to quantify the individual and combined effects of competition from the surrounding woody understory and ungulate browsing on long-term growth and survival of oaks. Our results show that, when it comes to survival, the relationship is relatively straightforward. Surrounding woody vegetation affects oaks’ survival in a negative direction, and so does browsing, even though the effect is smaller for the latter. When oaks experience both competition and browsing, the combined effect on survival is larger than either individual effect. For growth, however, the story becomes more complicated. Individually, both competition and browsing affects growth negatively, with browsing having the more substantial effect. When we analyze the combined effects, however, we find evidence suggesting that woody vegetation surrounding the oaks does not only have a competitive effect, but there are also elements of facilitation, through protecting the oak from browsing. Specifically, when we analyze the effects on oak height, the combined effect of competition and browsing has a smaller magnitude than the individual effect from browsing. The combined effect of competition and browsing on the basal diameter is, however, larger than either individual effect. Thus, the surrounding woody vegetation allows the plants to grow taller than if they only faced browsing, but this protection does not allow the plants to also grow in terms of basal diameter.

### Main contributions to the literature

The main contribution of the paper lies in the *joint* analysis of competition and browsing on long-term oak growth and survival. The previous literature has been able to identify individual negative effects from competition or browsing – with the latter literature being decidedly larger than the former – but so far, no one has been able to identify and quantify both the individual *and* the joint effects. As our results illustrate, explicitly analyzing the combined effects in addition to the individual ones allows us to gain insights that could not have been found by studying only the individual effects. For instance, we can show that when oaks face both competition from surrounding woody understory and ungulate browsing, the combined effect is smaller on height growth than if the oak had faced only browsing, even though competition and browsing individually both have negative effects on height growth. The conclusion that the surrounding woody understory therefore both competes with and facilitates the oak’s height growth could not have been drawn without our joint analysis.

In addition to identifying the joint effects of competition and browsing, our paper also makes an important contribution by quantifying the individual and combined effects. This allows us to compare the magnitude of various effects, and also provides an important background for practitioners wishing to learn how big an effect to expect from various changes in management strategy. Further, this also makes it possible to compare the competition and browsing effects between forest ecosystems.

### Implications for forestry and wildlife management

While the main aim of the paper is to shed light on the combined effects of competition and browsing on oak growth and survival, the rich data set collected in our large long-term field experiment also leads to interesting findings in other areas, often with a high practical management relevance. For instance, our results confirm what has been previously found in the literature, namely that if a forest manager is mainly interested in achieving fast regeneration of oaks, the best option from a growth as well as survival perspective is to protect the oaks with fencing, while also removing competing woody understory around the young oak trees. However, noting that fencing is not always an option – this could for instance be the case in forests with specific conservation value where fencing is considered too invasive – our results also suggest that protection by shrubs can be a viable alternative solution, in combination with controlling the surrounding woody vegetation to reduce the competition and reduction of the ungulate populations to reduce browsing.

Further, our findings also have direct application in wildlife management as our results show that fencing actually reduce the proportion of the tree species *Q. robur*, *P. tremula*, *S. caprea* and *S. aucuparia*. These tree species are not only highly palatable for ungulates but also act as important hosts for biodiversity. Our findings therefore suggests that managers of e.g. restoration and conservation efforts should not hesitate to allow browsing animals access to young plants in order to benefit the plants’ growing conditions, at least periodically and given that the local animal densities are not extremely high.

## Acknowledgement

We greatly appreciate Professor Frank Götmark’s efforts in maintaining the field experiment.

## Funding

The study was funded by the Swedish Research Council for the Environment, Agricultural Sciences and Spatial Planning (FORMAS), Linnæus University, Stiftelsen Extensus, Lars Hiertas Minne. and Stiftelsen Oscar och Lili Lamms minne.

## Author Contributions

Conceived and designed the experiments: AMJ, ML.

Collected the data: AMJ, LP, ML.

Analyzed the data: AMJ, MP, LP.

Wrote the paper: AMJ, MP, LP, ML, AF.

